# Improving Differential Expression and Survival Analyses with Sample Specific Compartment Deconvolution

**DOI:** 10.1101/2025.06.04.657908

**Authors:** Alisa Yurovsky, Richard Moffitt

## Abstract

**Motivation:** Studies on bulk RNA-seq of tumor biopsies can yield incorrect results because varying proportions of non-tumor tissues in the samples obscure the true signal and impact the accuracy of survival and differential expression analyses. Single-cell sequencing avoids these problems, but is still too expensive in clinical settings. Other deconvolution algorithms extract tissue-specific gene expression profiles from bulk sequencing, but cannot do this on a per-sample basis.

**Results:** We introduce SSCD - sample specific compartment deconvolution. SSCD extends non-negative matrix factorization with per-sample, per-gene constraint optimization. On simulated data, SSCD shows improvements in accuracy over existing methods. Using several real cancer datasets, we show that SSCD refines Differential Expression and survival analyses.

**Availability:** Code and data are available at https://github.com/ayurovsky/SSCD.

## Introduction

The advent of single cell sequencing (ssRNAseq) has greatly advanced the field of computational biology, enabling researchers to dissect cellular heterogeneity within complex tissues (Choi and Kim, 2019) and understand dynamic cellular processes (Cardona-Alberich et al., 2021). The ability to identify cell types via clustering pipelines like SINCERA (Guo et al., 2015) enables profiling of cell-type specific expression, answering critical questions of patient-to-patient variation.

According to recent reviews (Jovic et al., 2022; Wang et al., 2023; Nguyen et al., 2024), the primary limitation of single-cell sequencing remains its high cost. Aside from slowing the adoption of scRNAseq into clinical use, high cost limits the power of studies. Meanwhile, formalin-fixed paraffin-embedded (FFPE) samples are among the most widely available clinical specimens (Zhao et al., 2019), especially important for long-term rare disease studies. While technology is constantly improving, there remain many serious problems and challenges for scRNAseq on FFPE samples (Xu et al., 2023).

Bulk RNA sequencing (RNAseq) is significantly cheaper and more accessible, and remains a key technology for cancer genomics (Wang et al., 2020b; Thind et al., 2021). One of the main problems with RNAseq is the heterogeneous mixture of tissues/cell types present in the sample, illustrated in Fig. 1a. In the case of cancer, variation in tumor purity in biopsied samples greatly influence the clarity of the signal.

**Fig. 1.**
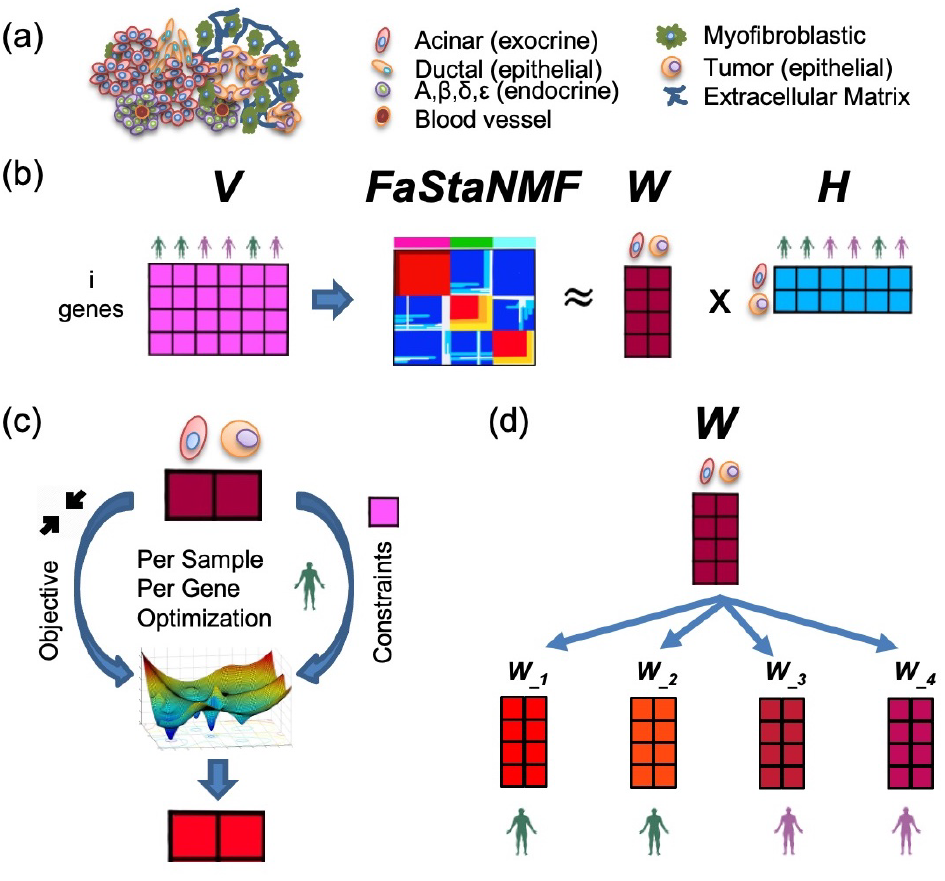
Description of SSCD, our optimization-based approach, where the goal is to obtain sample-specific and tissue-specific gene expression profiles from biopsies. **(a)** Patient biopsies are complex mixtures of cells in varying proportions, shown: pancreatic adenocarcinoma. **(b)** We take bulk RNA-seq expression data for all the samples (matrix V) and apply NMF deconvolution to obtain global tissue-specific expression profiles (matrix W) and proportions of tissues in each sample (matrix H). (C)We perform constraint optimization for each sample and each gene separately. **(d)** The final result is the deconvolution of tissue-specific expression profiles: an individual-specific matrix W i for every sample. These matrices can be used for differential expression studies and to identify patient-specific patterns in different tissues.

A sample with low tumor purity will obscure tumor-tissue-specific gene expressions, while a mixture with high tumor purity will obscure the signals from other tissues in the tumor microenvironment (TME).

Deconvolution is family of computational approaches used for extracting tissue specific profiles from RNAseq mixtures. In use for over a decade, deconvolution allows mining new tissue-specific insights from huge public repositories such as The Cancer Genome Atlas (TCGA) (Grossman et al., 2016) and Sequence Read Archive (SRA) Leinonen et al. (2011). There are many deconvolution methods which fall into two classes: reference-based approaches that require prior information, and reference-free composition-agnostic methods (Nguyen et al., 2024; Wang et al., 2020a). Many of the methods in both classes are based on non-negative matrix factorization (NMF) (Gaujoux and Seoighe, 2010), such as (Lin and Boutros, 2020; Qin et al., 2020; Tang et al., 2020). Reference-based methods include MuSiC (Wang et al., 2019) and CIBERSORT (Newman et al., 2019). Reference-free methods include CDSeq (Kang et al., 2019) and FaStaNMF (Sweeney et al., 2023).

Deconvolved bulk RNSeq expressions enable exploring tissue-level expression (ex. what is the average gene expressions profile for one subtype of specific cancer vs another). However, these global tissue expression profiles are not sample specific, and thus do not allow for differential expression or survival analyses *for specific tissues*.

We propose Sample Specific Compartment Deconvolution (SSCD), a new method extending FaStaNMF (Sweeney et al., 2023). Briefly, our method first applies FaStaNMF deconvolution to RNASeq samples, generating global tissue (compartment) expression profiles (Fig. 1b). We then perform per-sample per-gene optimization (Fig. 1c) to creates tissue-specific gene expression profiles for each sample (Fig. 1d).

Our approach opens opportunities for more detailed analysis for new and existing bulk-sequenced cohorts, including rare diseases and understudied populations. For example, a basic goal in a multitude of experiments is to find the difference between treatment and control groups. Very often this question is pertinent to either the tumor or the TME, and SSCD allows answering this question based on bulk RNAseq without bias from sample composition.

We first validate SSCD on synthetic datasets with known ground truth. We then apply SSCD to several cancer datasets. We recapitulate the results of a previous study of Pancreatic Adenocarcinoma (PAAD) (Moffitt et al., 2015), re-discovering tumor and stroma subtypes in a more straightforward manner. With data from the TCGA Adrenocortical Carcinoma study (Zheng et al., 2016), we demonstrate how SSCD-based analysis refines the stratification of patients with distinctive survival compared to a mixture-based approach. Finally, we look at cases before and after treatment (Nywening et al., 2016) to demonstrate the refinement of Differential Expression (DE) analysis.

## Materials and methods

### SSCD: Sample-Specific Compartment Deconvolution

The first step in our approach is to take the genes by samples matrix of mixture gene expressions (V in Fig. 1b), and deconvolve it with FaStaNMF (Sweeney et al., 2023). One result is the matrix H, where each row represents a tissue type (compartment), and the values are proportion for each sample. The second result is the matrix W, which provides the global gene expression profiles of deconvolved compartments. W and H post-processing follows the FaStaNMF methodology.

For each sample, for each gene we create an objective function and constraints and then perform optimization (Fig. 1c). R version 4.2.1 was used in this work. Optimization was performed with the alabama solver from the ROI package.

The objective is to minimize the differences between the global FaStaNMF-W and SSCD-W, using the sum of the relative square errors:

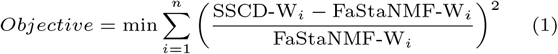

where n is the number of compartments. We enforce the non-negativity constraints using bounds. We set another hard constraint: SSCD values for each compartment will produce the original mixture values when multiplied by the FaStaNMF-derived H:

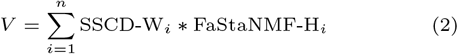

The above optimization process produces an new W matrix for each sample (Fig. 1d). The combination of the hard constraints and objectives allow the extraction of sample-specific differences for each gene, while preventing the solutions from wandering to far away from the global (average) tissue profiles.

### Signature gene selection for Sample-Specific Compartment Deconvolution

In well studied cases, we can use existing marker (or signature) genes from specific compartments for clustering and survival analysis. However, we need an automated method for general applications.

For tissue-agnostic signature gene selection, we consider each sample and each compartment individually. We select the genes that are high in this compartment, but low in other compartment(s). For each gene in compartment i:

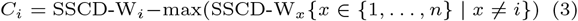

Then for each patient, for compartment i, we rank the genes. C_*i*_ values for the genes can be negative; the highest ranked genes are the most expressed in compartment i compared to other compartments. Then for each gene, we have all samples vote on the rank for compartment i by taking the median value across samples.

Top ranked genes are selected to become the signature genes for this compartment. This simple method is experiment and tissue agnostic, while allowing us to identify tissue specific signature genes.

### Datasets

#### Synthetic dataset construction methodology

In order to evaluate the ability of SSCD to reconstruct the individual tissue profiles from bulk RNASeq data, we need data with ground truth: exact tissue expression profiles for each sample and exact proportions of each tissue in each sample. We created two new synthetic datasets using the simulation methodology developed and introduced with FaStaNMF (Sweeney et al. 2023). Briefly, ground truth individual tissue expression profiles (W matrix) are created by random sampling and averaging from publicly-available single and bulk RNASeq data, while individual proportions of tissues are modeled with a Dirichlet distribution (H matrix). H and W ground truth matrices are then multiplied to create a synthetic matrix of bulk RNAseq gene expressions.

#### Resampled Single Cell (Synthetic)

Similarly to FaStaNMF paper (Sweeney et al. 2023), we obtained the data from the single cell transcriptome atlas of the human pancreas (Muraro et al. 2016) and created a dataset with 50 simulated patients and 12384 genes. The key differences in our dataset construction parameters stemmed from the desire to make the dataset a more challenging deconvolution problem (simulating four compartments), and more diverse for the purposes of showcasing sample-specific variation (sampling and averaging five cells per tissue per patient).

To create the H matrix, we set *α*_1_ = 0.4, *α*_2_ = 0.3, *α*_3_ = 0.2, *α*_4_ = 0.1 (representing alpha, beta, ductal, and acinar cells), and modeled the individual H proportions for each simulated patient with Dirichlet distribution. To make the W matrix for each sample, we randomly sampled and averaged 5 cells of each tissue types, and creating individualized tissue expression profiles. Then for each sample j, the mixed expression matrix V_*j*_ is calculated, where V_*j*_ = W_*j*_ H_*j*_.

Average tissue expressions for the dataset were obtained to demonstrate the complexity of the dataset. Supplementary Fig. S1a shows that the four tissues have low to moderate pairwise correlations, making it an easier deconvolution problem.

#### Gtex Resampled (Synthetic)

We obtained tpm-normalized bulk RNASeq gene expressions of four types of tissue (liver, lung, brain, and ovary) from healthy patients from the GTEx Portal (GTEx Analysis V8, downloaded on 06/21/2023). We filtered out uniformly-low expressed genes, keeping 13617 genes with mean >= 10 in at least one of the tissues. We then created a dataset of 50 simulated patients using the same methodology as above.

To create the H matrix, we set *α*_1_ = 0.4, *α*_2_ = 0.3, *α*_3_ = 0.2, *α*_4_ = 0.1 (representing liver, lung, brain, and ovary tissu and modeled the individual H proportions for each simulated patient with Dirichlet distribution. To make the W matrix for each simulated sample, we randomly sampled and averaged 40 real samples of each tissue types, and creating individualized tissue expression profiles (resulting in less individual variation than in the previous dataset). Then for each sample j, the mixed expression matrix V_*j*_ is calculated, where V_*j*_ = W_*j*_ H_*j*_.

These tissues have high pairwise correlations in comparison to the other synthetic dataset (Supplementary Fig. S1b), making it a harder deconvolution problem.

#### TCGA PAAD

We obtained clinical data and gene expression quantifications for Pancreatic Adenocarcinoma (PAAD) dataset from the

TCGA Research Network: https://www.cancer.gov/tcga. We retained 150 patient samples after filtering for whitelisted samples. Filtering out genes with zero expression across all samples, we retained 19986 genes.

#### TCGA ACC

We obtained clinical data and gene expression quantifications for Adrenocortical Carcinoma (ACC) dataset from the TCGA Research Network: https://www.cancer.gov/tcga. Filtering out patients without expression data, we retained 79 patient samples. Filtering out non-protein coding genes and genes with zero expression across all samples, we retained 19228 genes.

#### Linehan PAAD

We used a dataset of patients with locally advanced Pancreatic Ductal Adenocarcinoma (PDAC) from a phase Ib trial (Nywening et al., 2016) which were treated with FOLFIRINOX (majority of the patients were concurrently treated with CCR2 inhibitor PF-04136309). Enrolled patients had no prior treatment, underwent biopsies prior to the start of therapy and after treatment, which were then sequenced. We obtained processed gene expressions for before and after treatment samples from PurIST (Rashid et al., 2020) supplement (GSE131050), with a total of 66 samples. Filtering out non-protein coding genes and genes with zero expression across all samples, we retained 21421 genes.

## Results

### Validation on Synthetic Datasets

To assess how much the deconvolved values deviate from the truth we use Per Sample Average Absolute Deviation for true zero gene values:

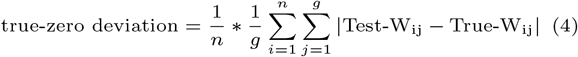

where n is the number of compartments and g is the number of genes. Similarly, we use Per Sample Average Relative Deviation for true non-zero gene values:

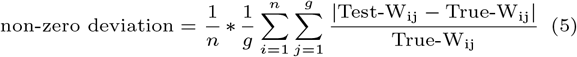

We need two separate functions because we want to assess the ability to both identify zero expression and estimate non-zero expression. This is especially important for the single cell-based simulation data.

This is a rough evaluation strategy, because it considers all genes, including highly expressed housekeeping genes that are not related to the “interesting” tissue-specific difference we focus on. i.e. we combine the deviations across all genes, whereas we know it is only the signature/marker genes that make the difference in downstream analyses. However, we want to show that we are doing no worse than other methods across all genes. Since no other approaches perform sample specific compartment deconvolution on bulk RNAseq data, we compare ourselves against three reference-free methods: a single run of NMF (from a standard R library), FaStaNMF (set for 200 iterations), and CDSeq (with default settings).

We repeated the experiments for the NMF-based methods three times (using different random seeds). For CDSeq, since we could not pass the random seed into the method, we used three different random seeds for sampling when creating the synthetic datasets. CDSeq estimated expressions were not on the same scale as the ground truth data, so we brought them into the correct range by scaling the mean of mixture values to the mean of CDSeq compartment results, separately for each compartment.

In Fig. 2, the violin plots show the distributions of errors for four methods on reconstructing gene expressions for two synthetic datasets. Fig. 2a shows the errors for the zero-expressed genes in the Resampled Single Cell dataset. SSCD has a smaller error distribution than FastaNMF 200, with the paired t-test p-value < 2.2e *-* 16. This is also the only case where CDSeq has a smaller error than SSCD. Aside from this exception, CDSeq performs the worst in all other cases compared to SSCD and FastaNMF 200, including on the zero-expressed genes in Gtex Resampled (Fig. 2c). Fig. 2b shows the errors for the non-zero expressed gene in Resampled Single Cell dataset, where SSCD has a smaller error distribution than FastaNMF 200, with the paired t-test p-value < 2.2e *-* 16. Fig. 2c shows the errors for the zero-expressed genes in the Gtex Resampled dataset; here, distribution of errors from SSCD and FastaNMF 200 are not statistically different, with the paired t-test p-value of 0.153. Finally, Fig. 2d shows a small (but statistically significant) improvement in the distribution of errors for non-zero expressed genes in the Gtex Resampled dataset (paired t-test p-value < 2.2e *-* 16).

**Fig. 2.**
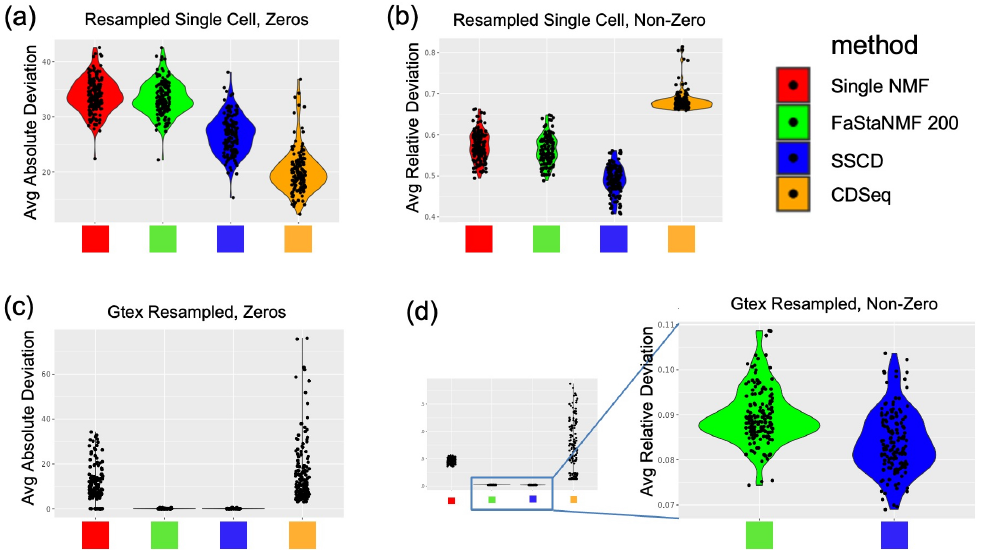
Distributions of errors for four methods on reconstructing gene expressions for two synthetic datasets. Violin plots in **(a)** and **(c)** show the distributions of errors for genes with zero expression, each dot representing the error averaged across all zero-expressed genes in a sample. Violin plots in **(b)** and **(d)** show the respective errors for all genes with non-zero expression, averaged per sample. In all cases, SSCD has either a statistically significant smaller error than FastaNMF 200 **(a, b, d)**, or shows no significant difference **(c)**.

These results demonstrate that SSCD performs as well as or better than FastaNMF 200 on the simulated datasets. Supplementary Fig. S2 makes a similar point in showing the true vs the reconstructed values for the four methods. The benefits of our method, however, lie in the extraction of the tissue-specific signal based on few marker genes, as we demonstrate in the following sections.

### Unsupervised Learning Application with TCGA PAAD

With real data, ground truth is not known. One application of SSCD is unsupervised learning for new subtype discovery, which we validate by recapitulating findings originally established on microarray data(Moffitt et al., 2015) in the RNAseq data from TCGA PAAD (Cancer Genome Atlas Research Network., 2017).

We performed a 3-compartment deconvolution on the bulk RNAseq data from TCGA. Based on the top 20 genes expressed in the global compartment profiles, we identified compartment 1 as Stroma, compartment 2 as Tumor, and compartment 3 as Normal pancreas. For marker genes, we used the 50 Stroma genes (25 normal and 25 activated) and the 50 Tumor genes (25 classical and 25 basal) from Moffitt et al. For each of the Stroma and Tumor compartments, we used the derived SSCD sample-specific compartment values of marker genes to perform consensus clustering on the samples.

In Fig. 3 panels a and c, survival difference between two patient groups (consensus clustering, k=2) recapitulate the outcome differences for two different stroma types found by (Moffitt et al., 2015). In Fig. 3 panels b and d, we show the survival difference between two main patient groups (consensus clustering, k=4, small clusters 3 and 4 discarded) for the tumor compartment. While the signal is weaker, we still see differences for two tumor subtypes re-identified in the TCGA PAAD cohort.

**Fig. 3.**
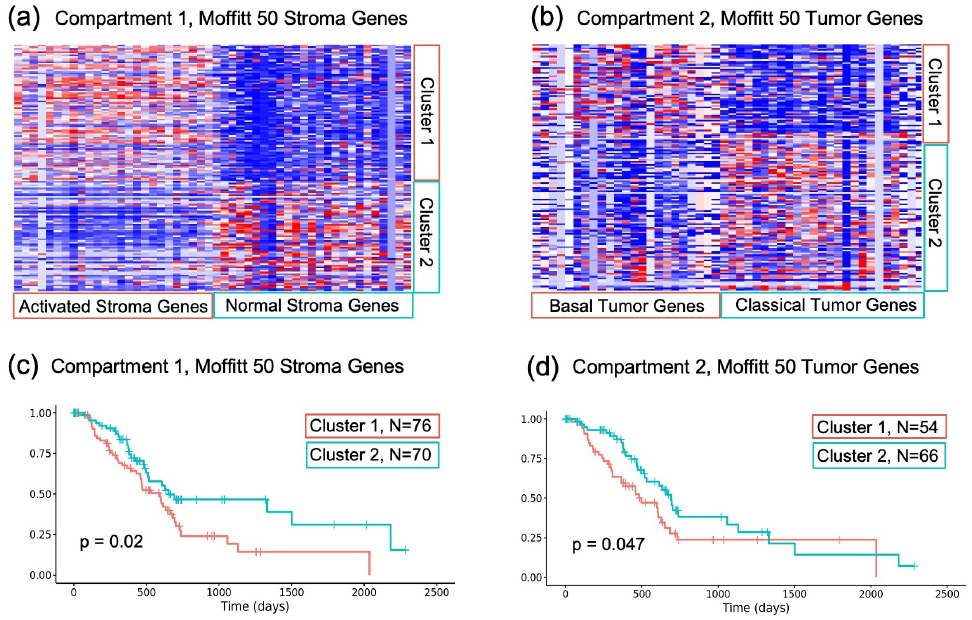
Clustering of TCGA-PAAD patients on SSCD-deconvolved Stroma compartment expressions for Moffitt 50 Stroma genes recapitulates significant survival differences for two Stroma sub-types **(a)** and **(c)**. Clustering the same patients on SSCD-deconvolved Tumor compartment expressions for Moffitt 50 Tumor genes has a weaker signal but still recapitulates survival differences for two Tumor sub-types **(b)** and **(d)**.

We furthermore compared the signature genes learned from the 3 SSCD-deconvolved compartments from the TCGA PAAD data (RNAseq: primary tumors) to those derived from the original study (Microarray: primary and metastatic samples). 50 SSCD Stroma genes had an intersection of size 15 with 50 Moffitt stroma genes, 50 SSCD tumor genes had an intersection of size 8 with 50 Moffitt Tumor genes, and 50 SSCD normal genes had an intersection of size 18 with 50 Moffitt Normal genes. For all three tissues, the p-value (Chi-square test) for the intersection between Moffitt list and signature gene list is the lowest possible in R (2.2e-16).

### Refining Survival Analysis with TCGA ACC

In this application, we demonstrate the ability of SSCD-based analysis to refine the stratification of patients into survival categories compared to a mixture-based method, using data from the TCGA Adrenocortical Carcinoma (ACC) study (Zheng et al., 2016). We also explore the use of our automatically selected signature genes.

Based on the two ACC tissue types (Weiss et al., 1989; Zheng et al., 2016), we performed 2 compartment deconvolution on the bulk RNAseq data. Using the top 50 variable genes from the mixture, we identified compartment 1 as tumor-steroid, and compartment 2 as tumor-immune. We concentrate on the tumor-immune compartment 2, as it is showing differences in survival. We used our signature gene selection to identify 50 marker genes from compartment 2.

In Fig. 4, panels a and d show the analysis using the bulk mixture expressions, with 50 top variable genes used for consensus clustering (k=2); two patient groups are identified with statistically significant survival differences. A hypothetical situation of using bulk mixture expressions with compartment 2 marker genes used for consensus clustering (Fig. 4 panels b and e) does not yield a statistically significant result, while the patient clustering is identical to the mixture. The intersection between 50 top variable genes and compartment 2 marker genes is 27, and thus the similar expression patterns are expected.

**Fig. 4.**
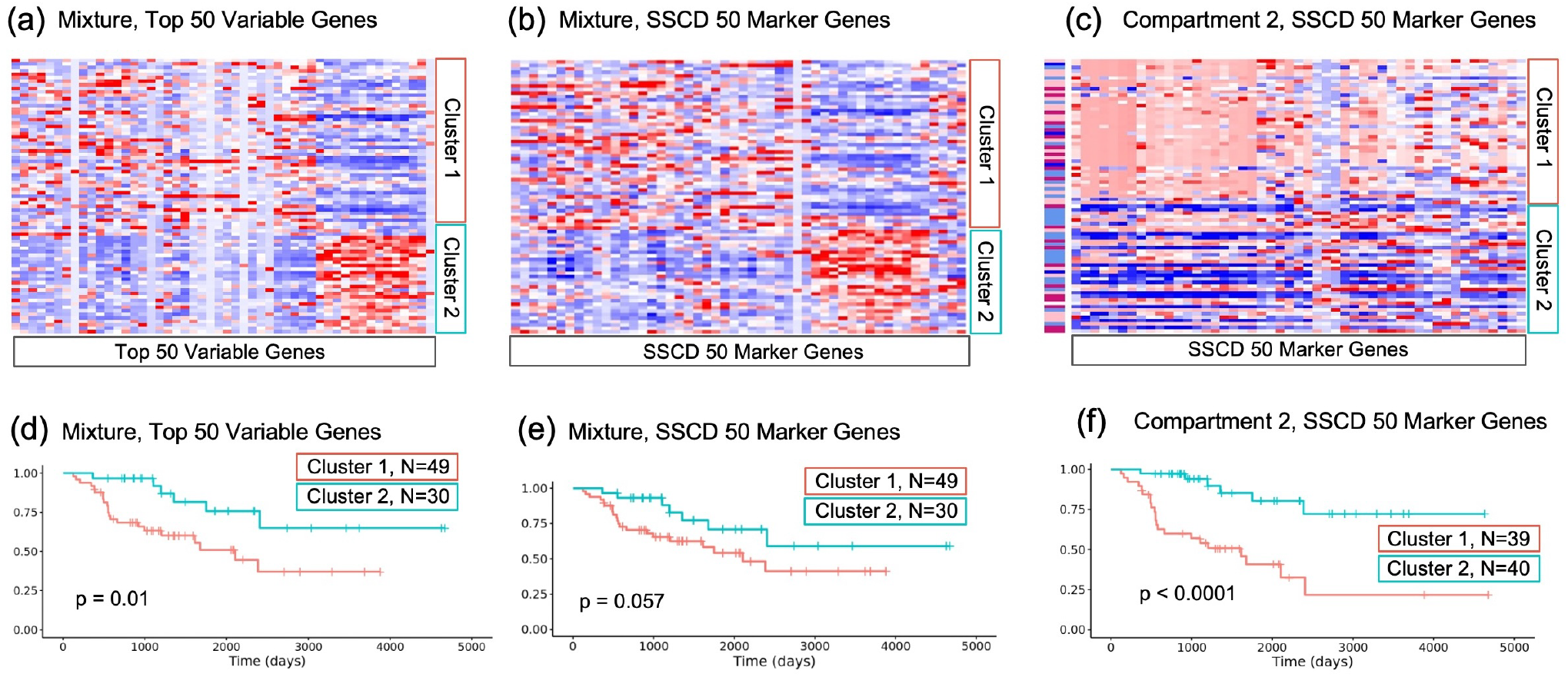
**(a)** Consensus clustering on mixture expressions with 50 top variable genes shows survival differences in **(d). (b)** Consensus clustering on mixture expressions using SSCD 50 marker genes does not show a statistically significant survival differences in **(e). (c)** However, consensus clustering on compartment 2 expressions using SSCD 50 marker genes gives a better separation with a lower p-value than **(a)** and 12 patients classified in a different survival category in **(f)**. Cluster assignment for compartment 2 shows low positive correlation (0.17) with low steroid phenotypes, a combination of two expression subtypes (“steroid-phenotype-low”(light blue) and “steroid-phenotype-low+proliferation”(dark blue)) found in (Zheng et al., 2016).

Finally, Fig. 4 panels c and f show compartment 2 isolated expressions with compartment 2 marker genes used for consensus clustering. The main conclusion from Fig. 4 is that SSCD-based survival analysis does indeed refine the mixture-based results (panel (d) vs panel (f)), with a lower p-value and 12 patients classified in a different survival category.

### Refining Differential Expression Analysis with Linehan PAAD

This application of SSCD focuses on improving the accuracy of DE analysis with another PAAD study by (Nywening et al., 2016). The study contained patient tumor biopsy samples before and after treatment with FOLFIRINOX and CCR2 inhibitor, and we we are interested in assaying the direct effects of tumor-suppressive treatment on the tumor gene expressions.

Similarly to TCGA PAAD, we performed 3 compartment deconvolution on the bulk RNAseq data. Based on the top 20 genes expressed in the global compartment profiles, we identified compartment 1 as immune-dominated Stroma, compartment 2 as Tumor, and compartment 3 as Normal tissues. The focus of our analysis is the tumor compartment. Applying SSCD to a before/after treatment PAAD patients’ dataset, we find that differential expression analysis on bulk RNA-seq mixture reveals incorrectly up-or-down regulated genes as compared to analysis on SSCD-extracted tumor compartment.

*Crucially*, Mixture-based analysis *incorrectly* identifies LDOC1 as one of the top 20 significant down-regulated genes for this experiment (log2FoldChange=-1.33, p-value=0.0207). On the other hand, Tumor-specific DE expression analysis does not find a statistically significant change for this gene. LDOC1 is known to be down-regulated in numerous cancers, including pancreatic (Nagasaki et al., 2003). The expected result after treatment is up-regulation or no effect.

*Equally importantly*, LRRC15 is associated with up-regulation in cancer progression in various studies, including pancreatic (Dominguez et al.), and the expected result after treatment is down-regulation or no effect; Tumor-specific DE expression analysis does not find a statistically significant change for this gene. However, Mixture-based analysis *incorrectly* identifies LRRC15 as one the top 20 significant up-regulated genes for this experiment (log2FoldChange=1.89, p-value=0.0214).

In Fig. 5 a, we explore the intersections between all significant DE genes from Mixture-based analysis, all significant DE genes from Tumor-specific analysis, and all genes that have a high correlation (absolute Spearman-rank correlation greater than 0.4) between mixture expressions and the H-matrix (tissue-specific abundance). We focus on the 141 genes that were uncovered with SSCD-enabled Tumor-specific analysis, which are enriched for genes associated with signaling pathways and linked to tumors. The full list is in Supplementary Fig. S3, while Fig. 5 b-c showcase some of these genes. For example, it is known that inhibition of MAP2K2 produces strong inhibition of pancreatic tumor cell growth, (Papademetrio et al., 2016), which is reflected in the decreased expression after treatment. The key insight from these figures is that while both Tumor-compartment-specific expressions and Mixture expression go down (or up) concordantly after treatment, SSCD enables capturing the small differences in sample-specific compartment-specific expressions, allowing these genes to be identified as deferentially expressed.

**Fig. 5.**
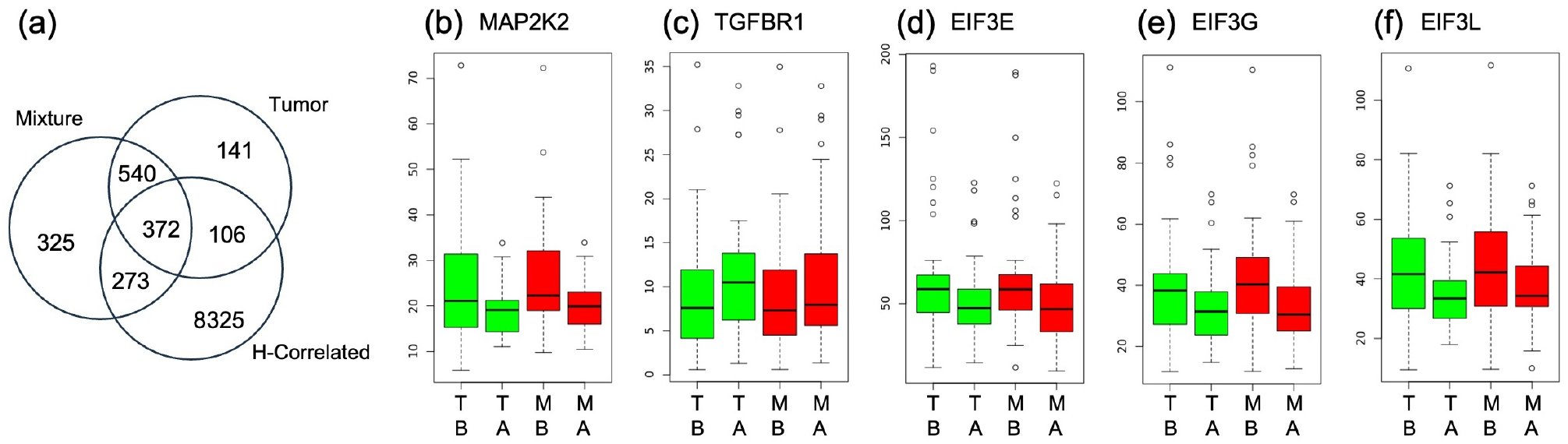
(a) Intersections between all significant DE genes from Mixture-based analysis, all significant DE genes from Tumor-specific analysis, and all genes that have a high correlation between mixture expressions and the H-matrix. (b)-(e) Expressions of several genes identified as significantly DE only in SSCD-identified tumor compartment. Green bars show the distribution of samples’ tumor compartment (T) gene expressions, before (B) and after (A) treatment. Red bars show the distribution of before and after bulk mixture (M) expressions.

Meanwhile, there is a significant overlap (273 genes) between significant DE genes from Mixture-based analysis and H-correlated genes, shown in Fig. 5 a. These genes are enriched for KEGG metabolic pathways, and do not appear to be tumor-related. The full list is shown in Supplementary Fig. S4.

## Discussion

While scRNAseq has greatly advanced the field of biology, the limiting factors of cost and technical problems are still relevant. On the other hand, bulk RNAseq obscures tissue-specific gene expressions. In this paper, we described Sample Specific Compartment Deconvolution (SSCD), a new method that uses bulk RNASeq data to create sample-specific, tissue-specific gene expression profiles. Through three different applications on cancer datasets, we demonstrated the utility of using SSCD for unsupervised subtype discovery, as well as refining survival and differential expression analyses.

The applications of SSCD show it to be useful for separability of the mixture in variable purity tumor samples. Basically, SSCD is useful whenever a single cell experiment could have been done, but it was too expensive, or bulk RNAseq (or formaldehyde-preserved) samples already exist. This experiment validates the use of SSCD for new subtype discovery by re-discovering the two tumor and two stroma subtypes found by Moffitt et al., but without the need for uniquely expressed genes (Moffitt et al., 2015). This type of analysis does not care about purity of individual tumor samples, so the clustering is less influenced by abundance, as in the original paper.

There are several future directions for SSCD. The first one would be to incorporate the principle component analysis (PCA) into SSCD. Our current optimization function works on a per sample per gene level and pushes the values together for all the compartments to minimize the error. We can extract latent compartment-specific information with PCA by taking the first principle component of each compartment for each sample. Then, each gene expression that is sample-specific *and* compartment-specific can be further refined after solving system of linear equations that incorporate the principle components. Another direction would be improve the speed of SSCD by rewriting the code in a compiled language. Finally, given the rapid expansion of the field of spatial transcriptomics, we would like to extend our deconvolution method to this domain, following the example of multiple reference-based and reference-free deconvolution methods (Li et al., 2023; Zhang et al., 2023).

## Data Availability

Code and data are at https://github.com/ayurovsky/SSCD.

## Acknowledgments

This work was supported by CRA CI and Stony Brook University IDEA Postdoctoral Fellowships.

