## Supplementary Figures for "Improving Differential Expression and Survival Analyses with Sample Specific Compartment Deconvolution"

(a)

| | acinar | alpha | beta | duct |
|---|---|---|---|---|
| acinar | 1.000 | 0.234 | 0.424 | 0.516 |
| alpha | 0.234 | 1.000 | 0.535 | 0.408 |
| beta | 0.424 | 0.535 | 1.000 | 0.791 |
| duct | 0.516 | 0.408 | 0.791 | 1.000 |

(b)

| | liver | lung | brain | ovary |
|---|---|---|---|---|
| liver | 1.000 | 0.872 | 0.911 | 0.787 |
| lung | 0.872 | 1.000 | 0.909 | 0.878 |
| brain | 0.911 | 0.909 | 1.000 | 0.824 |
| ovary | 0.787 | 0.878 | 0.824 | 1.000 |

**Supplementary Fig. S 1.** (a) Pairwise correlations for the four cell types in Resampled Single Cell dataset. Pairwise correlations for the four tissues in Gtex Resampled dataset.

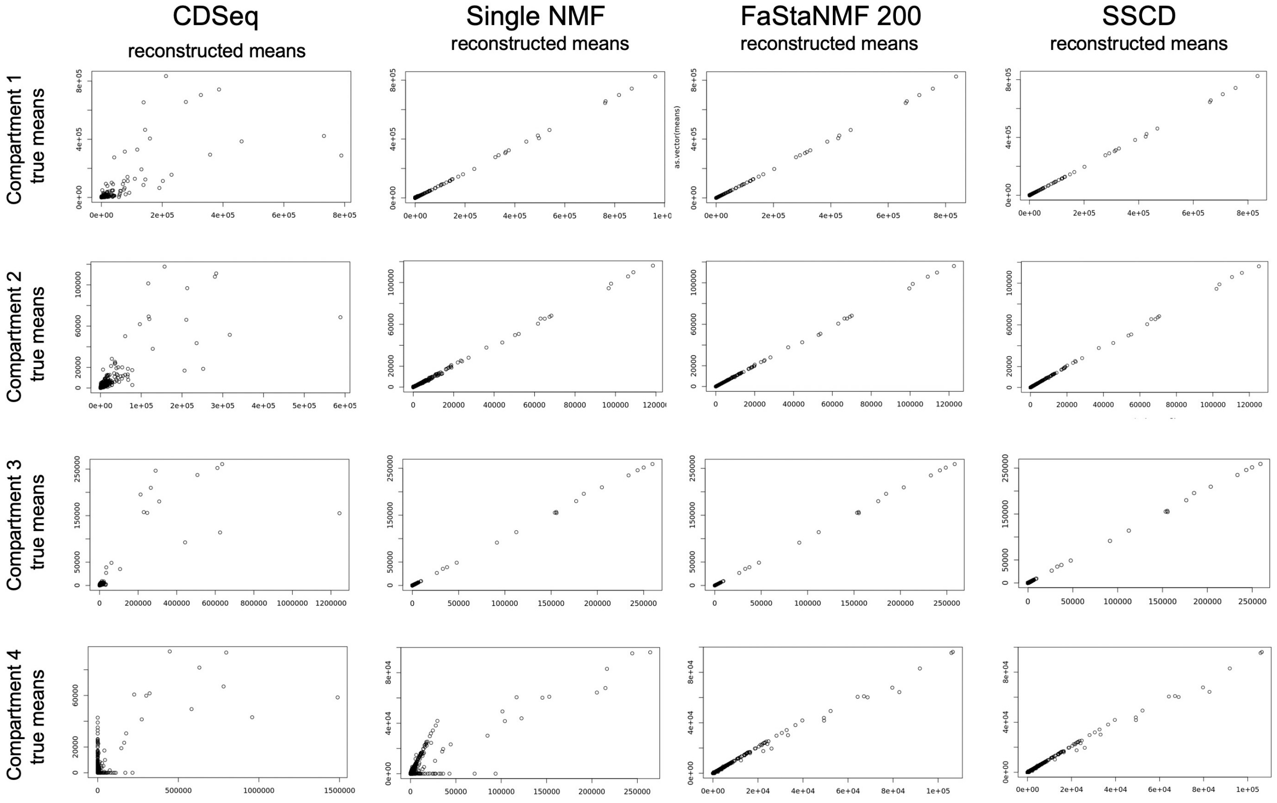


**Supplementary Fig. S 2.** True vs the reconstructed values mean values for the Resampled Gtex datasets for the four methods, stratified by the four compartments (tissue types).

AAED1, ABTB1, ADAMTS9-AS2, ADCY3, ALDH1A3, ALOX15B, ANP32B, ARL6IP5, ASNS, BGLAP, BICC1, C11orf45, C2CD4B, CA11, CA2, CD44, CDK14, CES1, CHST3, CIRBP, CLEC4D, CLMP, CNTN1, CORIN, CTDNEP1, CTRB2, CX3CL1, CXCL14, CXCR1, CXCR2, DCBLD2, DCHS1, DCXR, DGKZ, DKK3, DNAJC4, DPM3, EEF2, EIF3E, EIF3G, EIF3L, EML1, FAIM, FAM108A1, FAM92A1, FAT4, FERMT2, FILIP1L, FLJ42351, FSTL1, GGT3P, GJA1, GLT8D2, GNG10, GPX3, GTF3A, HCG27, HDAC9, HIGD2A, HIST2H2AB, HPGD, HPR, HSD11B1, HTRA1, HTRA3, IMPA2, ISM1, KAL1, KCND2, KLHL5, KMO, KRT23, LEPREL2, LGALS9C, LHFP, LINC00515, LINC00623, LMCD1, LMOD1, LOC728392, LRRC17, LTBP1, MAMDC2, MAP2K2, MIA2, MLLT3, MOXD1, MRPL53, MZT2B, NACA, NID2, NKD1, NPR3, NUAK1, OLFML3, OSR1, P2RX7, PARP9, PCDH18, PCOLCE, PDGFRB, PDZRN3, PI4K2A, PLEKHG4, PLG, POLR1D, PRICKLE1, PTPRM, RBMS1, RNASE1, SAMD9L, SDC2, SEC11C, SFMBT2, SLC2A4RG, SLC6A8, SNED1, SPSB3, SPTSSB, SRM, SSBP4, TGFBR1, THY1, TMEM160, TMEM255A, TMEM86A, TNFRSF25, TOP1MT, TPT1, TRABD2A, TREML2, TSPAN17, TWSG1, U2AF1L4, UBALD2, UBXN1, UFC1, UPF3A, UXT, VPS51, WDR18

**Supplementary Fig. S 3.** 141 genes that were uncovered with SSCD-enabled Tumor-specific DE analysis.

ABCC13, ABCC4, ABCD2, ABO, ACTL6A, ACYP1, ADAM15, ADAT1, ADIPOR1, AHSP, ALAS2, ALS2CR12, ANK1, ANKRD36BP2, AP1M2, AP2A1, APOL1, APOL2, ARG1, ARHGEF16, ASCC2, ASPHD1, ATF3, ATP5J2, BAIAP2L1, BCS1L, BLVRB, BNIP3L, BPGM, BZW2, C3orf37, C5orf4, C9orf78, CA1, CA12, CAPN13, CCDC176, CCNK, CCNL2, CDC20, CDC34, CENPM, CFL1, CGN, CHMP1A, CIB1, CKS1B, CKS2, CLK1, COX5B, CPT1B, CR1L, CRNDE, CSDA, CSNK2B, DCAF12, DCTN4, DDR1, DDX39A, DNAJB11, DNAJC6, DNTTIP1, DOK4, DPM2, DPY19L2P2, E2F2, EGLN3, EIF2AK1, EIF3I, EIF4A2, ELOVL1, EMC3, EMR1, ENO2, EPB41, EPB42, EPB49, EPS8L3, ERBB2, ERF, EXOC2, EZR, FAM104A, FAM117A, FAM210B, FAM46C, FBXO7, FECH, FKBP8, FLJ23867, FOXO3, FOXO4, FXYD3, GABARAPL2, GATA1, GINS2, GLRX5, GMPR, GPR146, GPR35, GUK1, GYPA, GYPB, GYPC, HAGH, HBA1, HBB, HBD, HBG1, HBG2, HBM, HBQ1, HDAC1, HDAC3, HEMGN, HIF1A-AS2, HINT1, HIVEP2, HMGA1, HMGCS1, HSBP1, IFIT1B, IFNK, IP6K2, ISCA1, JAZF1, KAT2B, KCTD20, KEL, KIAA1522, KLC3, KLF1, KRT1, LAMTOR5, LDHA, LINC00035, LINC00570, LOC91948, LPIN3, LYL1, MACF1, MAP2K3, MARCH2, MARCH8, MAT2A, MBNL3, MCM7, MDH2, METTL17, MFSD2B, MGST3, MKRN1, MLF2, MORC3, MPP1, MRPS21, MRPS5, MXI1, MYL4, NADSYN1, NCOA4, NDUFB4, NDUFB8, NFE2, NFKB2, NOXA1, NPRL3, NSDHL, NXF1, OAZ1, OR2W3, OSBP2, P2RX5, P4HA1, PAGE2B, PCGF5, PDCD6, PFKP, PGRMC1, PHOSPHO1, PIH1D1, PIM3, PITHD1, POLE2, PPFIA4, PPIA, PPM1G, PPP4C, PPP5C, PRDX6, PRELID1, PROSER1, PRSS8, PSMD8, PSMF1, PSORS1C3, PTK6, QARS, R3HDM4, RAB2B, RANBP10, RBM3, RBM38, RBM4, RGS10, RHAG, RHOB, RNF10, RNF11, RUNDC3A, S100A6, SELENBP1, SEMA3B, SEMA4B, SESN3, SFN, SH2D3A, SIAH2, SLC25A37, SLC25A39, SLC29A2, SLC2A1, SLC41A3, SLC4A1, SLFN14, SLX1A, SLX1B, SMIM1, SNCA, SORD, SOX6, SPIRE2, SPPL2B, SPTA1, SPTB, SRRD, SSB, SSU72, ST6GAL2, ST6GALNAC1, ST6GALNAC4, ST7-OT4, STRADB, STX16, TAF1C, TAL1, TBCEL, TCP11L2, TMBIM1, TMC4, TMCC2, TMED9, TMEM214, TMEM92, TMOD1, TMPRSS4, TNS1, TRAPPC2L, TRIM38, TRIM58, TSPAN5, UBB, UBE2O, UBXN6, UPK1A-AS1, VRK1, WDR67, WHAMMP3, YOD1, YPEL3, ZC3H12A, ZC3HAV1, ZER1

**Supplementary Fig. S 4.** Overlap of 273 genes between significant DE genes from Mixture-based analysis and H-correlated genes.
